# The lack of trade-off between conformational stability and binding affinity in a nanobody with therapeutic potential for a misfolding disease

**DOI:** 10.1101/2024.09.15.612864

**Authors:** Atanasio Gómez-Mulas, Athi N. Naganathan, Angel L. Pey

## Abstract

To improve protein pharmaceuticals, we need to balance protein stability and binding affinity with *in vivo* efficiency. We have recently developed a nanobody (NB-AGT-2) against the alanine:glyoxylate aminotransferase with high stability (*T*_m_∼85°C) that may be useful to treat a misfolding disease called primary hyperoxaluria type 1. In this work, we characterize the relationships between protein stability and binding affinity in NB-AGT-2 by generating single and double cavity-creating mutants in its hydrophobic core. These mutations decrease thermal stability by 10-20 °C, reflecting changes in thermodynamic stability of up to 8 kcal·mol^-1^, hardly affecting their binding affinity for its target. Our results thus show that NB stability can be challenged without an effect on its binding.

## Introduction

Nanobodies (NBs) are single-domain proteins derived from the heavy-chain antibodies of camelids, with a great potential as therapeutic agents [1–7]. NBs display high conformational stability, with an unfolding free energy in the range of 10-20 kcal·mol^-1^ and well-described by a two-state equilibrium model of unfolding [8,9]. Their thermal stability is often high, i.e. in the range of 70-90°C midpoint melting temperature. The interaction of target with an NB is mediated by three hypervariable regions (typically loops) called the Complementarity-Determining Regions (CDR 1-3). The CDR 3 is likely the most important CDR for binding to its target and it appears to contribute substantially to NB stability [1,10]. However, the relationships between the conformational stability of NB, their binding affinity for their target and their potential as therapeutics are not well understood.

We have recently generated NB against the human alanine:glyoxylate aminotransferase 1 (AGT-1). Alterations in AGT-1 (e.g. by mutations) cause a deficiency to detoxify glyoxylate and generate an overproduction of oxalate that causes renal failure [11]. The NBs previously generated (named as NB-AGT) showed high conformational stability and high affinity for AGT (dissociation constants, *K*_d_, from nM to pM, [9]).

In this work, we have used NB-AGT-2 as a model to study structure-stability-function relationships in a therapeutic NB. NB-AGT-2 shows a high stability (∼ 20 kcal·mol^-1^ at room temperature by chemical denaturation) and ∼ 86 °C (∼ 359 K) of midpoint denaturation temperature (*T*_m_). It also binds tightly to AGT-1 (with a *K*_d_ value around 0.3 nM). Therefore, NB-AGT-2 is an excellent model to study structure-stability-function relationships for therapeutic NBs. We have carried out these studies by generating single and double mutants at two buried positions of hydrophobic residues (L22 and I72) to create cavities (mutations to V and A) in the protein core of NB-AGT-2. These cavities largely reduce the thermodynamic stability of the NB-AGT-2 at room temperature without affecting its binding affinity, in contrast to a previous study in which *humanizing* mutations in NB often increased thermodynamic stability with a concomitant large decrease in binding affinity [12]. Thus, our experimental stability and binding studies together with thermodynamic statistical mechanical calculations provide novel insights into the structure-function-stability relationships of NB as therapeutic agents.

## Materials and methods

### Protein expression and purification

*E. coli* BL21 (DE3) cells were transformed with the pET-24(+) vector containing the cDNA of NB-AGT-2 WT (wild-type) and several variants at L22 and I72 positions carrying a C-terminal 6His-tag. Variants were generated chemically by GenScript (Leiden, The Netherlands) with codons optimized for expression in *E. coli*. A preculture of 240 mL of LB medium containing 30 µg·mL^-1^ kanamycin (LBK) was inoculated with transformed cells and grown for 16 h at 37 ºC. These cultures were diluted into 4.8 L of LBK, grown at 37 ºC for 3 h and nanobody expression was induced by the addition of 0.5 mM IPTG (isopropyl β-D-1-thiogalactopyranoside) and lasted for 6 h at 25 ºC. Cells were harvested by centrifugation and frozen at -80 ºC for 16 h. Cells were resuspended in binding buffer, BB (20 mM Na-phosphate, 300 mM NaCl, 50 mM imidazole, pH 7.4) plus 1 mM PMSF (phenylmethylsulfonyl fluoride) and sonicated in an ice bath. These extracts were centrifuged (20000 g, 30 min, 4 °C) and the supernatants were loaded into IMAC (immobilized-metal affinity chromatography, Cytiva, Barcelona, Spain) columns, washed with 40 bed volumes of BB and eluted in BB containing a final imidazole concentration of 500 mM. These eluates were immediately buffer exchanged using PD-10 columns (Cytiva, Barcelona, Spain) to 50 mM HEPES(N-2-Hydroxyethylpiperazine-N □-2-ethanesulfonic Acid)-KOH pH 7.4 and stored at -80 °C upon flash freezing in N_2_. NB-AGT samples were further purified by loading them onto a SuperDex 75 10/30 size exclusion chromatography column (Cytiva, Barcelona, Spain) using 20 mM HEPES-NaOH, 200 mM NaCl pH 7.4 as mobile phase at 0.5 mL·min^-1^ flow rate. Fractions containing NB-AGTs were collected, concentrated, buffered exchange to 50 mM K-phosphate pH 7.4 and stored at -80 ºC after flash-freezing in liquid N_2_. Purity and molecular weight were analysed again by SDS-PAGE (dodecyl-sulphate polyacrylamide gel electrophoresis). NB-AGT-2 concentration was measured using the molar extinction coefficients (*ε*_280_=33015 M^-1^·cm^-1^) according to their primary sequence.

### Differential scanning calorimetry (DSC)

DSC experiments were carried out using a VP-DSC differential scanning microcalorimeter (Malvern Pananalytical, Malvern, UK) with a cell volume of 137 µL and automated sampling. Experiments were performed in 50 mM K-phosphate using 20 µM of NB-AGTs and a scan rate of 2 °C·min^-1^. Scans were carried out in a temperature range of 20-100 °C to allow complete unfolding and minimize the effects from irreversible thermal denaturation. In fact, the reversibility is quite significant although not complete, thus supporting the applicability of equilibrium denaturation to determine the relevant unfolding parameters [8,9,12] (reversibility of 40±10% for all nine variants under these experimental conditions).

To evaluate the thermal denaturation behavior of NB-AGTs, we applied a model in which denaturation was reversible but not presumed to be a two-state process (i.e. only the native and unfolded are populated). To this end, we used a simple approach in which the temperature (*T* in K) dependence of the apparent heat capacity (*C*_p,app_) was described by equation 1:

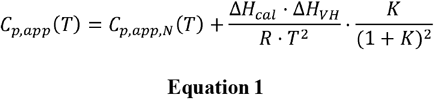

Where *C*_p,app,N_ is the temperature-dependent apparent heat capacity of the native state (described by a straight line equal to a+b·*T*), the area under the calorimetric *peak* or calorimetric enthalpy (Δ*H*_cal_), the Van’t Hoff enthalpy (Δ*H*_VH_). Please note that two different temperature *T* scales along the manuscript, one in °C for a broader readership and one in K for some thermodynamic calculations (e.g. Equations 1-2). Both scales are equivalent for differences in *T*_m_.

### Isothermal denaturation by Guanidium Hydrochloride (GdmHCl)

GdmHCl denaturation of variants of NB-AGT-2 were carried out by mixing protein solutions with different concentrations of GdmHCl (typically in the 0-6 M range, GdmHCl concentrations were determined using refractive index measurements). Experiments were carried out in 50 mM K-phosphate pH 7.4 using 5 µM of NB-AGT-2 variants. Samples were incubated at 4 °C for 24 h then the temperature was increased stepwise (18, 25, 32, 39 and 46 °C) remaining at each temperature for 20 min before measurements. Protein unfolding was monitored by fluorescence spectroscopy at each temperature, using an excitation wavelength of 295 nm and emission in the range 320-380 nm (both with 5 nm slits). Blanks in the absence of protein were routinely measured and subtracted. Spectroscopic measurements were carried out using quartz 3 × 3 mm cuvettes. To monitor denaturation, we used the ratio of emission intensities at 365 and 335 nm (I_365_/I_335_).

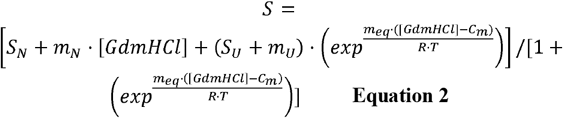

where *S* is the experimental spectral feature (I_365_/I_335_) as a function of the [GdmHCl], *S*_N_ and *S*_U_ are the fitted spectral features for the Native and Unfolded state baselines at 0 M GdmHCl, respectively, *m*_N_ and *m*_U_ are the slopes of the native and unfolded state baselines, *m*_eq_ describes the unfolding cooperativity, *R* is the ideal gas constant and *T* is the experimental temperature (in K). This model provides an excellent description of chemical denaturation of NB-AGTs as well as other NBs [8]. The product of *C*_m_ and *m*_eq_ provides an estimation of the unfolding Gibbs free energy (Δ*G*_UNF_).

### Surface Plasmon Resonance (SPR)

Binding affinity for the interaction between NB-AGT-2 variants and the AGT-1-LM variant was evaluated using a Biacore T200 Surface Plasmon Resonance instrument (Cytiva, USA). All six NB-AGTs were individually and covalently immobilized on NTA chips aiming for 500 response units (RUs). For this, we first determined the amount of time required for each variant to reach this level of response. To capture the His-tagged NB-AGTs, the nitrilotriacetic acid (NTA) chip was loaded using a NiCl_2_ 0.5 M solution and 100 - 200 nM nanobody samples in a HBS-P+ buffer (10 mM HEPES, 150 mM NaCl, 0.05% v/v Tween20) passed under a flow of 5 µL·min^-1^. Once these times were determined, the immobilization procedure required activation of the NTA matrix with the same 0.5 M NiCl_2_ solution and activation of the carboxyl groups of the same matrix with a 1:1 mixture of N-ethyl-N-(3-diethylamino-propyl)-carbodiimide (EDC) and N-hydroxysuccinimide (NHS). Nanobodies followed for the time previously determined under a 5 µL·min^-1^ flow and ethanolamine blocked the remaining activated carboxyl groups, ending the immobilization step. Each NTA chip has 4 flow cells in which, the first was activated with the EDC/NHS mixture and then blocked with ethanolamine without ligand ever flowing through so as to function as a reference for the others. Experiments were carried out at 25 °C.

Affinity constants (*K*_a_; their inverse are the dissociation constants, *K*_d_) as well as the dissociation and association rate constants (*k*_on_ and *k*_off_, respectively) were determined by performing Sigle Cycle Kinetics (SCK) assays. In this case, serial dilutions of the different AGT variants in HBS-EP+ (10 mM HEPES, 150 mM NaCl, 3 mM EDTA, 0.05% v/v Tween20), with concentrations ranging 0.74-60 nM, were injected in 120 s pulses under a 70 µL·min^-1^ flow. Resulting sensograms were fitted to a 1:1 interaction model and kinetic constants obtained using the Biacore T200 Evaluation Software.

The changes in binding free energy (ΔG_binding_) between a variant and the WT protein were determined from the corresponding *K*_d_ values (*K*_d(variant)_ and *K*_d(WT)_) using the following equation: 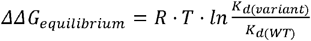 **(Equation 3)**, where R is the ideal gas constant (1.987 cal·mol^-1^·K^-1^) and *T* is 298.15 K. Similarly, the changes in activation free energies were determined for the association (ΔΔ*G*^□^_on-rate_) and dissociation reaction (ΔΔ^□^) from the rate constants (*k*_on_ and *k*_off_ values of the variants and the WT protein) from the following equations: In all the cases, the errors reported for these variables were those obtained from linear propagation of fitting errors.

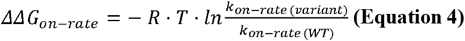

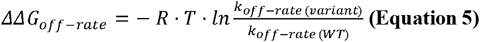

### Statistical mechanical analysis of the structural perturbation induced by mutations

The block Wako-Saitô-Muñoz-Eaton model (bWSME) [13,14]) was employed to understand the extent of mutational propagation using the structure of NB-AGT-2 (its structure was predicted from its sequence using AlphaFold2 [9,15]. Briefly, two consecutive residues are considered to fold-unfold in unison, enabling the partitioning of NB-AGT-2 variants into 59 blocks. Each of the blocks can be fully folded (binary variable *1*) or unfolded (*0*), and together with the single-sequence approximation (SSA), double-sequence approximation (DSA) and DSA with interaction across islands (DSAw/L) substates, the phase space of NB-AGT-2 is partitioned into 938,504 microstates. The statistical weight of each of the microstates is estimated by combining van der Waals packing interactions observed in the native structure (*E*_*vdW*_, determined by strength of van der Waals interactions per native contact ξ within a 5 Å heavy atom cut-off radius), all-to-all electrostatics (*E*_*elec*_), simplified solvation (Δ*G*_*solv*_, determined by the heat capacity change per native contact 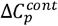), and a destabilizing entropic penalty term for fixing residues in the native conformation (Δ*s*_*conf*_). The resulting total partition function z is then employed to calculate free energy profiles as a function of the number of structured blocks, heat capacity profiles, positive coupling free energies (Δc_+_) and perturbation profiles (ΔΔ*G*_+_) [14]. The parameter ξ was alone tuned to reproduce the melting temperatures of the variants observed in experiments. The final parameters are: 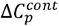 of -0.36 J mol^-1^ K^-1^ per native contact, ξ of -87, -84.3, -86.3, and -83.5 J mol^-1^ for the variants of NB-AGT-2 WT, L22A, I72A and the double mutant L22A-I72A, respectively, and Δ*s*_*conf*_ of -14.5, 0 and -20.6 J mol^-1^ K^-1^ per residue for ordered residues (helical or beta strand), proline and glycine/coil residues, respectively (as observed in the native structure). All simulations were performed assuming pH 7 protonation states for charged amino acids and at an ionic strength of 0.1 M. For the calculation of perturbation profiles (ΔΔ*G*_+_ = Δ*G*_+,*Mut*_ - Δ*G*_+,WT_), the Δc_+_was first calculated for the WT, mutation introduced via PyMol, and finally the altered contact map was fed into the model to estimate Δ*G*_+_ for the mutant.

## Results and discussion

### Selection of mutants and their expression

To select cavity-creating mutations that could perturb the structural integrity and dynamics of the protein core of NB-AGT-2, we first modeled its expected structure using Alphafold-2 (https://www.dnastar.com/software/nova-protein-modeling/novafold-ai-alphafold-2/?gclid=Cj0KCQiA6fafBhC1ARIsAIJjL8kYtxC-SweC55xlDFGLol8d_bH647CvsHQd9gGXCuYOHcPRleS3TAkaAiYUEALw_wcB) [9,15,16]. From the model with the highest scoring, we selected two fully buried positions in the protein core as calculated using GetArea [17](Figure 1) and containing hydrophobic and bulky residues (L22 and I72). L22 is located far from the CDR loops in this model (shorter distance > 15-20 Å from all three CDR) whereas I72 is at a distance of ∼6 Å from CDR2, and ∼15 and ∼12 Å from CDR3 and 1, respectively (Figure 1). To progressively perturb this core region, we introduced single and double mutations to valine and alanine. Due to the high stability of NB-AGT-2 (Table 1) [9], all the variants expressed well in *E*.*coli* and were amenable for biophysical characterization.

**Table 1.** Energetics of denaturation of NB-2 variants determined by DSC. Denaturation parameters are derived from fittings of a unfolding equilibrium model to the experimental data (Equation 1). Errors are those from fittings or derived from them by linear propagation.

| **NB-AGT-2**<br>**variant** | **$T_m$ (°C)** | **$\Delta H_{cal}$ (kcal·mol<sup>-1</sup>)** | **$\Delta T_m$ (°C)** |
| --- | --- | --- | --- |
| **WT** | 85.60±0.01 | 94.5±0.4 | 0.00±0.02 |
| **L22V** | 80.09±0.01 | 78.8±0.4 | -5.51±0.02 |
| **L22A** | 73.37±0.01 | 74.8±0.4 | -12.23±0.02 |
| **I72V** | 84.17±0.01 | 95.3±0.3 | -1.43±0.02 |
| **I72A** | 78.92±0.01 | 85.9±0.3 | -6.68±0.02 |
| **L22V-I72V** | 78.72±0.02 | 72.2±0.5 | -7.02±0.03 |
| **L22V-I72A** | 72.78±0.02 | 61.2±0.6 | -12.82±0.03 |
| **L22A-I72V** | 71.23±0.02 | 57.4±0.6 | -14.37±0.03 |
| **L22A-I72A** | 64.72±0.03 | 49.2±0.5 | -20.88±0.04 |

**Figure 1.**
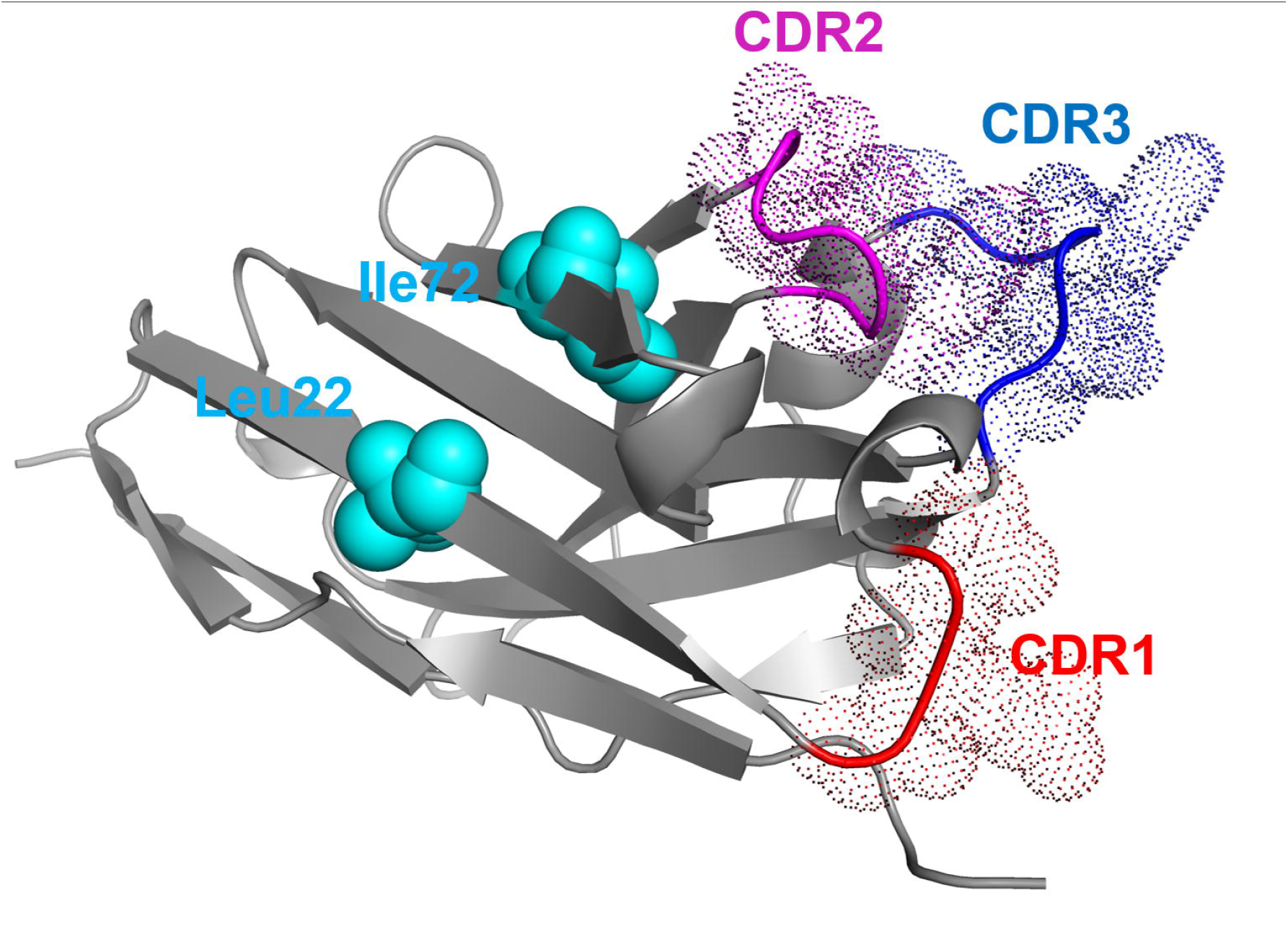
Hydrophobic residues selected for cavity-making using the model of NB-AGT-2 obtained by Alphafold 2 [9,15,31]. Two residues were selected as cavity-making (Leu22 and I72) due to their full burial using this model and the algorithm GetArea (https://curie.utmb.edu/getarea.html)[17] and their distance to the hypervariable loops in the CDR1-3 are also highlighted (CDR1.- 28-30; CDR2.- 54-59; CDR3.- 103-108).

### Gradual thermal destabilization upon cavity-making mutations

Upon protein purification, we first measured the thermal stability of the different variants by differential scanning calorimetry (DSC). The DSC scans were highly symmetric, and fits to a reversible unfolding model were good given the experimental conditions, with reversibilities of 40% (average of all nine NB-AGT-2 variants). The ratio between calorimetric and Van’t Hoff (Δ*H*_cal_ and Δ*H*_VH_, respectively) was overall 0.8±0.1, supporting that a simple two-state unfolding describes well the thermal denaturation of all NB variants [8,9,18].

We observed a gradual and clear destabilizing effect of the cavity-making mutations at L22 with weaker effects of mutations at L72 (Figure 2 and Table 1). It is clear that the changes in the half-denaturation temperature (*T*_m_ values) have additive effects on thermal stability for the double mutants at L22 and L72 (Table 1). The temperature-dependence of Δ*H*_cal_ resulted in a value of 2.3±0.3 kcal·mol^-1^·K^-1^ (Figure S1), consistent with theoretical calculations of the unfolding heat capacity change (Δ*C*_p_) for the two-state reversible unfolding of a protein of this size (1.8-1.9 kcal·mol^-1^·K^-1^)[19,20].

**Figure 2.**
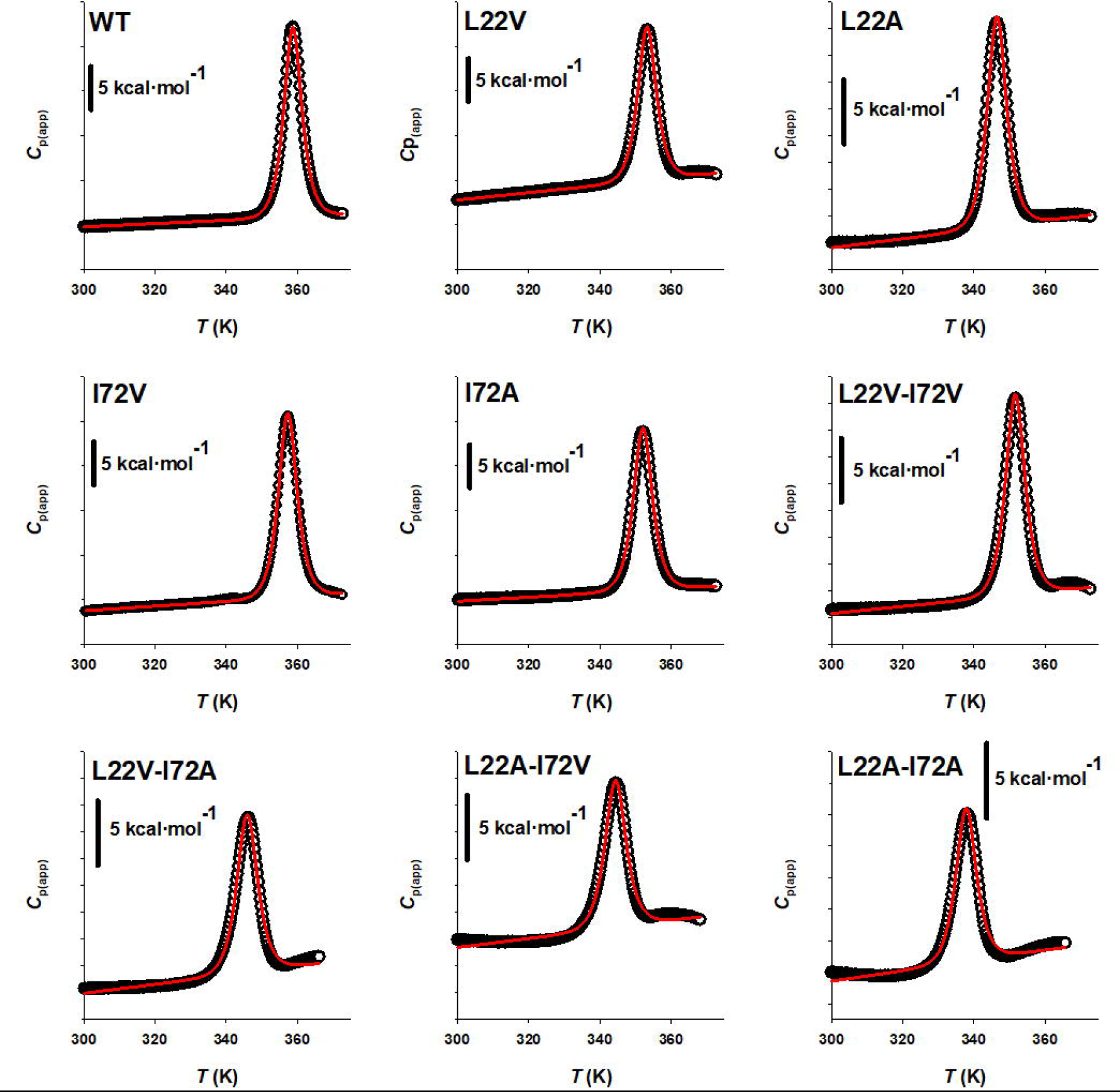
Differential scanning thermograms showing destabilization of NB-AGT-2 due to cavity-creating mutations. Closed symbols show experimental data and red lines are best fits to a reversible unfolding model (Equation 1). Energetic denaturation parameters can be found in Table 1. Please note that the x-axes are shown in absolute temperature (in K) for the purpose of fittings.

### Thermal destabilization reflects a lower thermodynamic stability caused by cavity-making mutations

Since thermal denaturation of NB-AGT-2 variants showed some degree of reversibility (even though in some cases the thermal scan ended at temperatures 20 ^°^C higher than the end of the calorimetric transition, see Figure 2), it is plausible that the effects on thermal stability reflect changes in thermodynamic stability at room temperature as seen for NB-AGT-2 WT [8,9]. To test this hypothesis, we have carried out denaturation experiments with GdmHCl as denaturant at temperatures much lower than the *T*_m_ of the variants (18-46 ^°^C) [9]. We selected the WT and four mutants of NB-AGT-2 showing thermal destabilization from moderate to large (Δ*T*_m_ between -5.5 to - 20.9 ^°^C, see Table 1). Representative GdmHCl denaturation profiles are shown in Figure 3A and were fitted well by a simple two-state equilibrium denaturation [8,9,12]. Using the average *m*_eq_ values for these five variants at a given temperature (see legend of Figure 3), we derived the unfolding Gibbs free energy changes (Δ*G*_UNF_) for all of them. Results at different temperatures provide a consistent picture of mutational effects on thermodynamic stabilities (Figure 3B), with a destabilization (from ΔΔ*G*_UNF_, average±s.d from five temperatures) ranging from -2.4±0.7 (in L22V) to -8.0±1.4 (in L22A-I72A) kcal·mol^-1^ (Figure 3B). Indeed, despite the small set of variants studied, the linear correlation between thermodynamic (ΔΔ*G*_UNF_) and thermal destabilization (Δ*T*_m_) is excellent (Figure 3C), with a r^2^ value of 0.97 and a slope of 0.36±0.05 kcal·mol^-1^ of chemical destabilization per 1°C of thermal destabilization. It is worth noting that upon using the Schellman equation [21] and the thermodynamic parameters *T*_m_ and Δ*H*_cal_ values, we obtain a dependence of 0.26 kcal·mol^-1^· °C^-1^, in good agreement with our analyses (Figure 3C). These results reinforce the notion that under our conditions kinetic distortions minimally affect thermal denaturation of all NB-AGT-2 variants studied, since similar results are obtained when thermodynamic analyses are carried out using reversible chemical unfolding and partially reversible thermal unfolding experiments [9].

**Figure 3.**
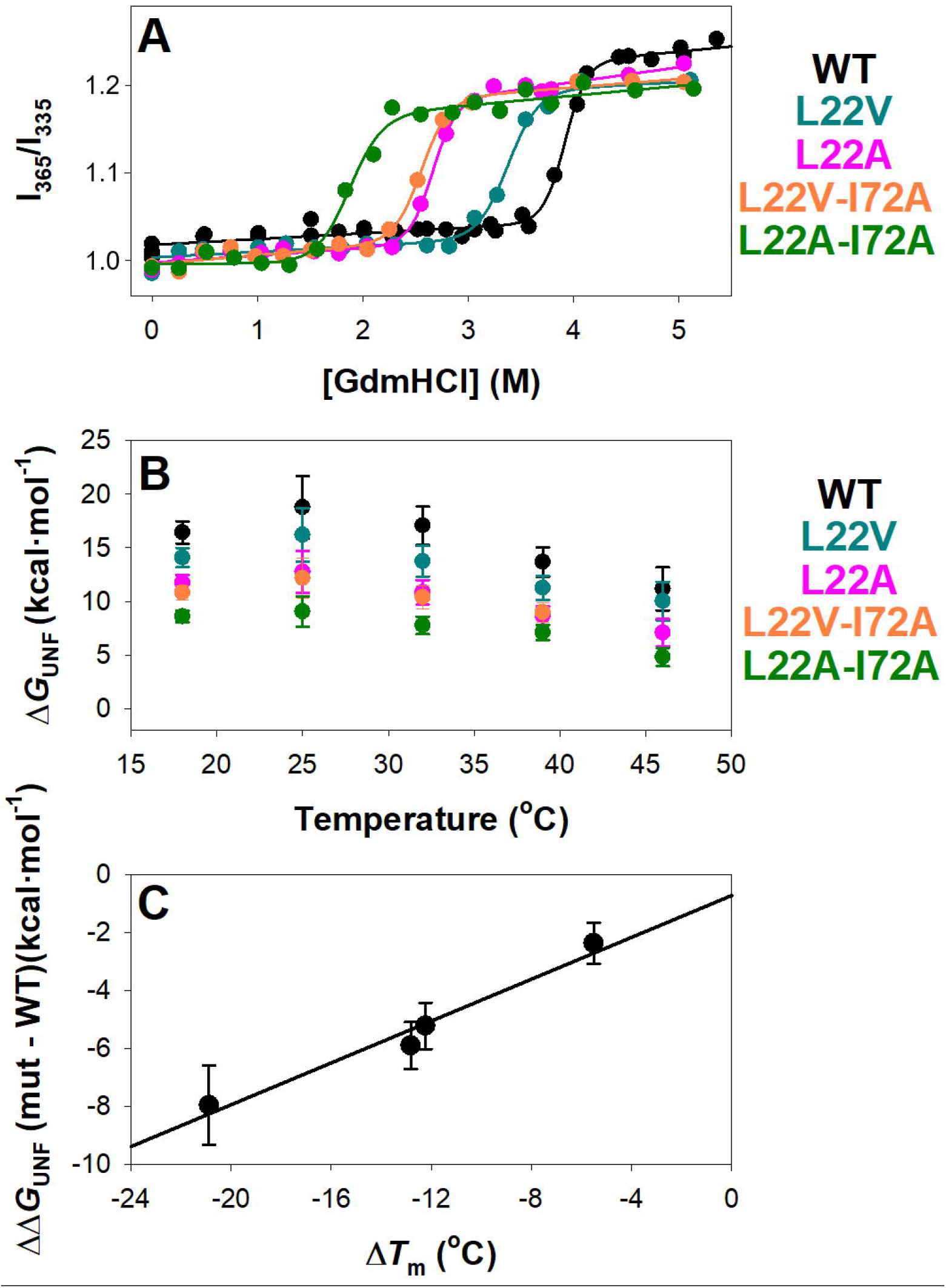
The effect of cavity-making mutations on thermal stability reflects the effects on thermodynamic stability at room temperature. A) Isothermal denaturation of NB-AGT-2 variants by GdmHCl followed by fluorescence at 25°C. B) Unfolding Gibbs free energy changes (Δ*G*_unf_) for GdmHCl isothermal denaturation in the temperature range 18-46 ^°^C. *m*_eq_ values used to calculate ΔG_UNF_ were the average of all variants at a given temperature: 18 ^°^C.- 4.13±0.26; 25 ^°^C.- 4.79±0.74; 32 ^°^C.- 4.29±0.45; 39 ^°^C.- 3.92±0.39; 46 ^°^C.- 3.70±0.68 (in kcal·mol^-1^·M^-1^). C) Linear correlation between thermodynamic destabilization (as the Δ*G*_UNF_ of a given variant and the WT protein, as the mean±s.d. at five temperatures, ΔΔ*G*_UNF_) and the thermal destabilization (as the different in *T*_m_ between a given mutant and the WT protein, ΔΔ*T*_m_). The r^2^ value is 0.97 and the slope is 0.36±0.05 kcal·mol^-1^·^°^C^-1^.

### Weak correlation between thermodynamic stability and binding affinity

To test whether thermodynamic stability of NB-AGT-2 would have functional implications, such as effects on the binding affinity for its target AGT-1-LM, we have carried out surface plasmon resonance (SPR) analyses of the interaction for the WT NB-AGT-2 and four variants that caused widely different thermodynamic destabilization (ranging from -2.4±0.7 in L22V to -8.0±1.4 in L22A-I72A, in kcal·mol^-1^). A summary of these results is reported in Figure 4. First, we must note that these four variants tested bound with similar affinities to the WT NB-AGT-2, with dissociation constants (*K*_d_) values in the 0.3-0.9 nM range (Figure 4B), yielding a maximal change in binding Gibbs free energy (ΔΔ*G*_binding_) due to mutations of ∼ 0.6 kcal·mol^-1^. This is much larger than the thermodynamic destabilization of the least stable variant (−8.0±1.4 kcal·mol^-1^). Consequently, when we plot the change in binding free energy (ΔΔ*G*_equilibrium_) as a function of the thermodynamic destabilization (ΔΔ*G*_UNF_), their correlation is weak (r^2^=0.64 and slope of -0.055±0.024)(Figure 4C). As we show in Figure 4B-C, the small changes in binding affinity arise from mild and opposing effects of the mutations on the *k*_on_ and *k*_off_ values, which are both slightly increased in the variants thus almost cancelling their effects on the *K*_d_ values (Figure 4B). Linear fits showed in Figure 4C provide slopes of 0.082±0.030 (ΔΔ*G*^□^_on-rate_, r^2^=0.72) and 0.136±0.049 (ΔΔ*G*^□^_off-rate_, r^2^=0.72) for the dependence of activation binding free energies on thermodynamic stability. These analyses support that the cavity-making mutation slightly destabilize the complex with AGT-LM, possibly due to perturbations of stabilizing interactions in the complex, but also somehow decrease the free energy barrier for the association with AGT-1-LM.

**Figure 4.**
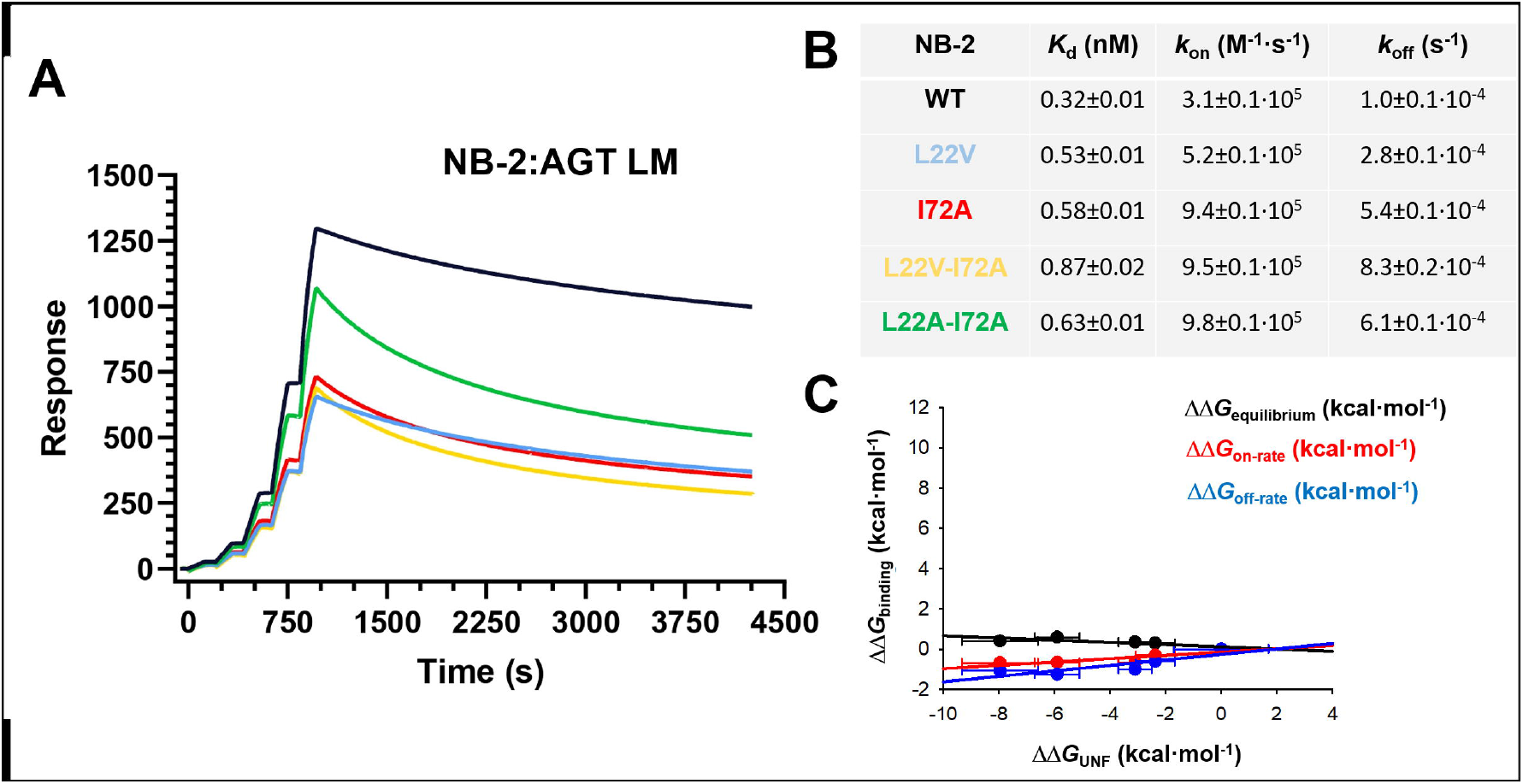
Binding of NB-AGT-2 variants to AGT-1-LM. A) Sensograms corresponding to the binding of NB-AGT-2 WT to AGT LM. NB-AGT-2 was covalently attached to the chip and AGT-1-LM at increasing concentrations were sequentially injected. B) Equilibrium dissociation constants (*K*_d_), as well as association (*k*_on_) and dissociation (*k*_off_) rate constants. C) Weak correlation between the effects of variations in NB-AGT-2 and binding free energies. ΔΔ*G*_equilibrium_ show values derived from *K*_d_ values of a variant and the WT NB, ΔΔ*G*^□^_on-rate_ show values from *k*_off_ values and ΔΔ*G*^□^_off-rate_ show values from *k*_on_ values. Errors are those from propagation using standard errors from fitting to a 1:1 binding model.

### Statistical mechanical analysis provide insight into mutational effects on the function and stability of NB-AGT-2

To provide further structural and energetic insights into the relation across stability, functionality of NB-AGT-2, and the extent of mutational effects, we employed the structure predicted by AlphaFold2 [15] (Figure 1) as a template for structure-based statistical mechanical calculations (the block Wako-Saitô-Muñoz-Eaton model (bWSME) [13,14]) (see Material and Methods, Statistical mechanical analysis of the structural perturbation induced by mutations). We carried out these calculations on NB-AGT-2 WT, two single mutants L22A and I72A, and their double mutant L22A-I72A (whose melting temperatures span over 20°C in thermal stability with respect to the WT variant)(Table 1).

The only input into the model are the predicted structure and experimental differences in stability. The latter is used to calibrate the model by modulating a single parameter as discussed in the Materials and Methods section. The resulting free energy profiles as a function of the natural model coordinate, the number of structured blocks, reveal a three-state-like behavior with the native state being the most populated (N) in the WT protein. On introducing destabilizing mutations, the population of the partially structured intermediate (I, particularly in case of the double mutant) and the unfolded state (U) increase (low values of the reaction coordinate in Figure 5A). It is possible that the lack of differences in the binding affinity could be a consequence of the minimal effect of mutations on CDR3, or it could still be a significant effect which is *rescued* by the large stability of the protein. To address this question, we quantified coupling free energy differences between the WT and the mutant which condense the microstate diversity, and energetic variability onto a single residue-level estimate shedding light on the extent of mutational perturbations. As we show in Figure 5B-C, the mutation L22A has pleiotropic effects, affecting the entire protein structure (residues as far as 20 Å from the mutated sites, an effect observed for highly disruptive mutations in other proteins [22–28], and this effect is enhanced in the double mutant L22A-I72A. On the other hand, the mutant I72A has a minimal effect on the overall coupling free energies and hence the native structure. Thus, though the consequence of mutations is varied and context-dependent, they translate to little overall effect on the final binding affinity possibly due to the large stability of the protein.

**Figure 5.**
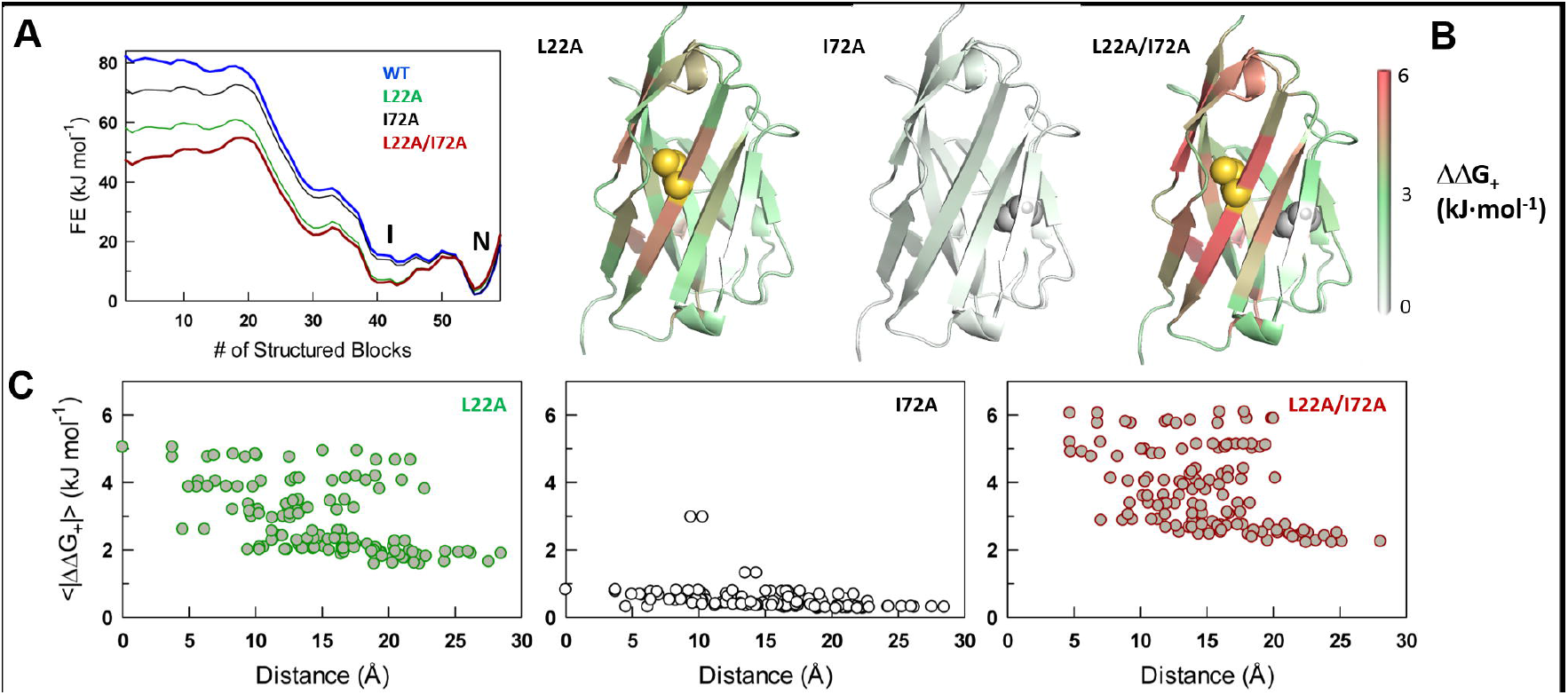
Statistical mechanical calculations reproduce mutational effects on the conformational ensemble of NB-AGT-2 variants. A) One-dimensional free energy profiles (FE) along the folding reaction coordinate. Note that cavity-creating mutations increasingly destabilize the native conformation (N) to both intermediate (I) and unfolded conformations. B) Differences in positive coupling free energies plotted onto the structural model of NB-AGT-2. The differences between selected mutants and the WT NB-AGT-2 are displayed according to the color scale. C) Same as panel B, but plotted as a function of C*α* -C*α* distance from the mutated site (in the case of the double mutant we used the average distance between the mutated sites).

## Conclusions

In protein engineering we must face a trade between protein stability and function [29]. This is particularly challenging when we are designing protein pharmaceuticals [30]. However, this trade-off is not always easy to be understood, and therefore, improvements for protein pharmaceuticals are not an easy task.

The potential of NB to treat human diseases is large. NB are single-domain antibodies naturally produced by camelids and derived from heavy-chain antibodies, and therefore, these are small proteins (∼15 kDa, about one-tenth the size of a conventional antibody) that retain the extreme affinity of conventional antibodies [1]. NB show excellent properties as pharmacological agents, since these are highly specific, stable (weeks to months in a fridge or at physiological temperature), low-immunogenicity and are relatively less expensive to generate. In this work, we have shown that the exceptional properties of a NB raised to treat a misfolding disease (PH1) can support several challenges. Decreasing their thermal stability by ∼20 degrees or their thermodynamic stability by ∼10 kcal·mol^-1^ at room temperature have little effects on their extreme affinity for their target. Though two very disruptive mutations (such as in the mutant L22A/I72A) are introduced in the protein core, the affinity for its target remains surprisingly unaltered. It is interesting to note that in an earlier publication [12], the authors generated a quadruple mutant (F/Y42V, E49G, R50L and G52W, named *VGLW*) in a highly conserved region (framework region 2, FR2) next to the CDR 1 and/or 2 in order to *humanize* it, and potentially make it more suitable for therapeutic applications. This quadruple mutant only showed local changes in the structure, increasing the NB stability (by ∼13 kcal·mol^-1^) but reducing the affinity for its target by ∼50-fold [12]. In the same work, using a different NB as template in which a 11-fold mutant with mutations humanizing the NB outside the FR2, the same quadruple mutant *VGLW* reduced by 200-fold the affinity for its target and causing pleiotropic changes in the structure (no data on stability was reported)[12]. Remarkably, we show that a single (L22A) and a double mutant (L22A-I22A) affect the stability of almost the entire structure with a concomitant decrease in protein stability but without detrimental effects on its binding affinity. Statistical mechanical modeling further suggests that mutational effects mostly stabilize non-natively folded states.

Overall, we conclude that structure-stability-function relationships in NBs are very complex. Our work thus provides novel insight into the fundamental thermodynamic and binding properties of NBs and show that this technology has the potential to be improved as protein pharmaceuticals.

## Abbreviations

AGT-1: alanine:glyoxylate aminotransferase 1
CDR: Complementarity-Determining Regions
*C*_m_: the midpoint of chemical denaturation
*C*_p,app_: apparent unfolding heat capacity
DSC: differential scanning calorimetry
HEPES: N-2-Hydroxyethylpiperazine-N□-2-ethanesulfonic Acid
*K*_a_: equilibrium association constant
*K*_d_: equilibrium dissociation constant
*k*_off_: dissociation rate constant
*k*_on_: association rate constant
NB: nanobody
NB-AGT-2: a nanobody generated to target AGT
*m*_eq_: the cooperativity index for chemical denaturation
RU: response units
SPR: surface plasmon resonance
*T*_m_: mid-point denaturation temperature
WT: wild-type
Δ*G*_BINDING_: equilibrium binding free energy change
Δ*G*_UNF_: unfolding free energy change
Δ*G*_+_: positive energy coupling
Δ*H*_cal_: calorimetric enthalpy for unfolding
Δ*H*_VH_: Van’t Hoff enthalpy for unfolding
ΔΔ*G*□_off- rate_: difference in binding activation free energy between a variant and the WT protein
ΔΔ*G*□ _on-rate_: difference in dissociation activation free energy between a variant and the WT protein
ΔΔ*G*_+_: difference in positive energy coupling between a variant and the WT.

## Contributions

A.L.P conceived the project. A.G-M carried out protein purifications and all experimental work. A.N.N. carried out statistical mechanical analysis. A.L.P drafted the manuscript and all the authors contributed to the final version.

## Acknowledgements

We thank Dr. Juan Luis Pacheco-Garcia for technical help. We also thank Prof. Eduardo Salido for insightful comments on the manuscript.

