## Supplementary Data for "The lack of trade-off between conformational stability and binding affinity in a nanobody with therapeutic potential for a misfolding disease"

**Supplementary Information**

**Figure S1. Temperature-dependence of unfolding enthalpies (Δ*H*_cal_) .** Black symbols are those determined experimentally, and the solid line assumes a linear dependence on temperature that yield a value for the unfolding heat capacity change (Δ*C*_p_).

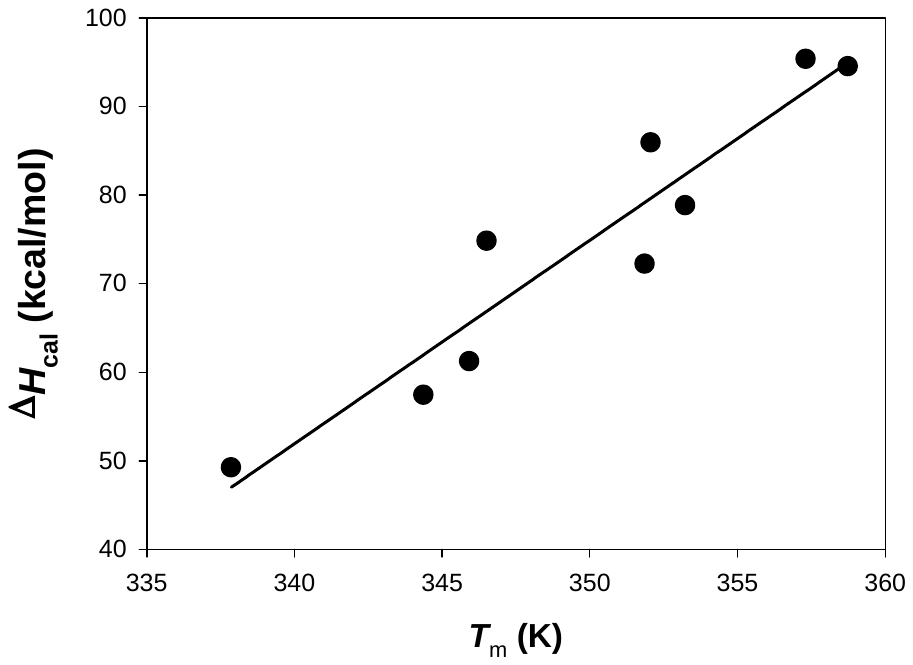
